# Towards resolution of the intron retention paradox in breast cancer

**DOI:** 10.1101/2022.03.04.482791

**Authors:** Jaynish S. Shah, Michael Milevskiy, Veronika Petrova, Amy YM Au, Justin J.L. Wong, Jane E. Visvader, Ulf Schmitz, John E.J. Rasko

## Abstract

After many years of neglect in the field of alternative splicing, the importance of intron retention (IR) in cancer has come into focus following landmark discoveries of aberrant IR patterns in cancer. Many solid and liquid tumours are associated with drastic increases in IR and such patterns have been pursued as both biomarkers and therapeutic targets. Paradoxically, breast cancer (BrCa) is the only tumour type in which IR is reduced compared to adjacent normal breast tissue.

In this study, we have conducted a pan-cancer analysis of IR with emphasis on BrCa and its subtypes. We explored mechanisms that could cause aberrant and pathological IR and clarified why normal breast tissue has unusually high IR.

Strikingly, we found that reduced IR in BrCa can be largely attributed to normal breast tissue having the highest occurrence of IR events compared to other healthy tissues. Our analyses suggest that low numbers of IR events in breast tumours are associated with poor prognosis, particularly in the luminal B subtype. Interestingly, we found that IR frequencies negatively correlate with cell proliferation in BrCa cells, i.e. rapidly dividing tumour cells have the lowest number of IR events. Aberrant RNA binding protein (RBP) expression and changes in tissue composition are among the causes of low IR in BrCa.

Our results suggest that IR should be considered for therapeutic manipulation in BrCa patients with aberrantly low IR levels and that further work is needed to understand the cause and impact of high IR in other tumour types.

## Introduction

Pre-mRNA splicing is a ubiquitous process that is crucial for the maintenance of transcriptomic complexity and gene expression regulation in eukaryotic cells^1,2^. Perturbations to this highly calibrated system can have severe consequences and lead to diseases including cancer^3–6^. In this context, numerous studies describing intron retention (IR) in disease have shed light on the mechanisms leading to aberrant and pathological IR^7–9^.

The importance of IR in cancer has been emphasised following landmark discoveries about (i) aberrant IR patterns in leukemia^10,11^, (ii) IR as a source of neoepitopes^12^, (iii) tumour suppressor gene inactivation by intronic polyadenylation^13^, (iv) IR-based biomarkers^14,15^, and (v) IR as a therapeutic target^16^.

IR is regulated by *cis*- and *trans*-acting modulators^2,17,18^ facilitating cellular responses to a range of environmental stimuli^19^. Intron-retaining mRNA transcripts are often degraded via nonsense-mediated decay (NMD), thereby causing down-regulation of the host gene. The burden of IR in disease is governed by perturbations to mechanisms known to regulate this form of alternative splicing, including mutations, splicing factor dysregulation, and epigenetic variations.

However, despite the cumulative evidence for the importance of IR in cancer, a systematic analysis of IR regulation in BrCa and the role of aberrant IR in BrCa biology has not been conducted to date. In this study we sought to resolve the paradox wherein breast cancer exhibits reduced IR, which is an important consequence of alternative splicing.

We analysed 615 BrCa patient transcriptomes which included four major molecular subtypes (Luminal A, Luminal B, Basal, and Her2 positive). We confirmed a consistent downregulation of IR in BrCa. However, we also observed that normal breast tissue has a significantly higher IR event frequency compared to other healthy tissues. The number of IR events correlated with survival in the luminal B BrCa subtype. Differences in IR frequencies are largely influenced by the tissue’s cellular composition as well as specific dysregulated RNA binding proteins (RBPs).

## Results

### High IR is associated with improved survival in Luminal B subtype breast cancer

To compare IR profiles across human cancers we retrieved transcriptomics data for nine different solid tumours and matched adjacent normal tissues from The Cancer Genome Atlas (TCGA; Figure 1A) and quantified IR using the IRFinder algorithm, which we have previously validated^17^. Overall, we identified a total of 11,943 unique IR events (Supplementary Table 1), of which 917 were shared among all nine cancers analysed.

**Figure 1.**
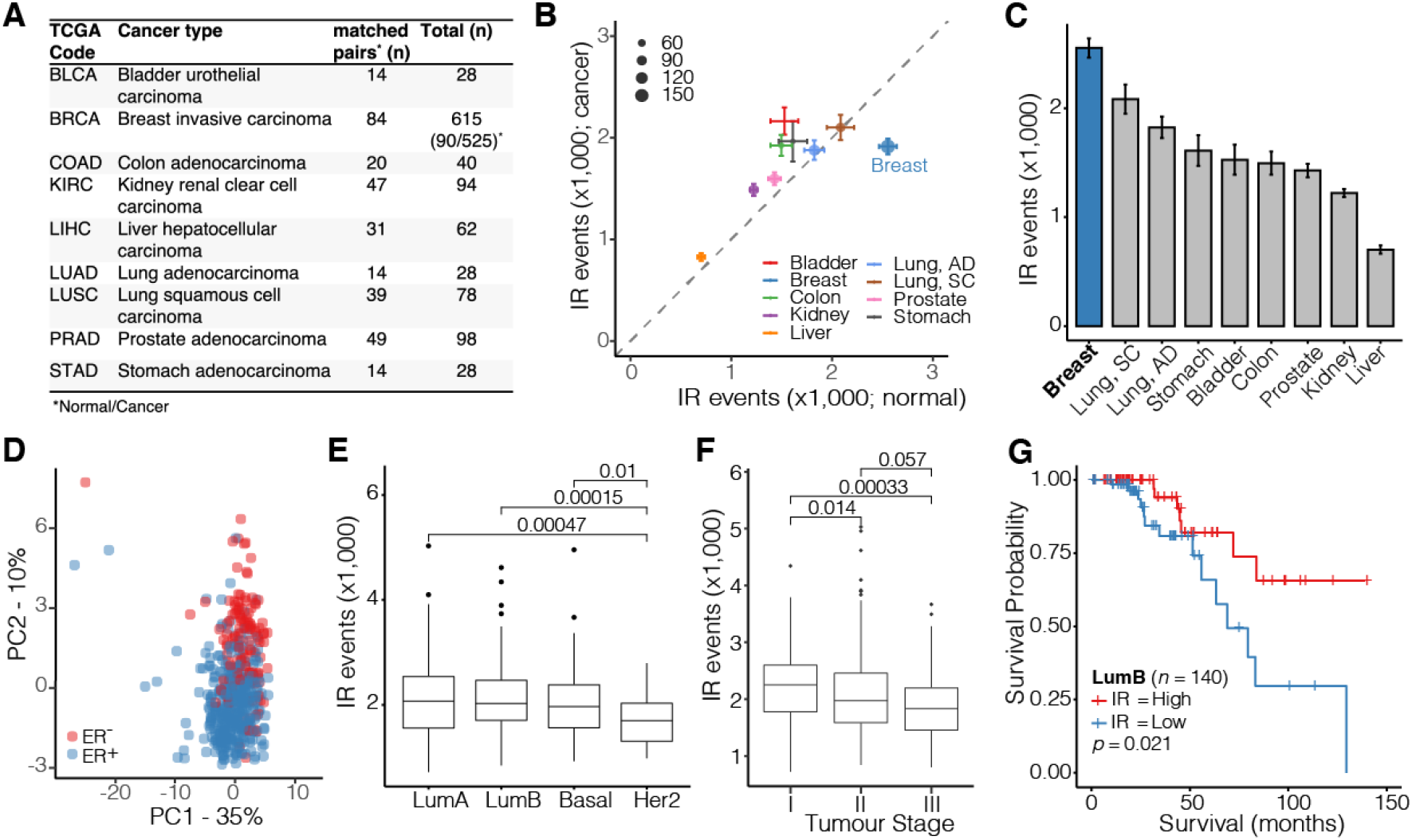
Breast cancer intron retention in the TCGA cohort. **(A)** Overview of TCGA samples analysed. **(B)** Scatter plot illustrating the average number of IR events in cancers vs adjacent normal tissues. Error bars indicate standard deviations. The size of the dots is proportional to the number of samples analysed. **(C)** Bar plot showing the average number of IR events in normal tissues in descending order. Error bars represent standard error of the mean. **(D)** PCA plot showing clusters of specific IR profiles in ER positive and ER negative tumour samples (*n* = 509). **(E)** Distributions of IR event frequencies in four major BrCa subtypes (LumA – Luminal A; LumB –Luminal B; Basal, HER2 - human epidermal growth factor receptor 2 positive). **(F)** Distributions of IR event frequencies in tumour samples assigned to three tumour stages. **(G)** Kaplan-Meier plot illustrating the survival probabilities of Luminal B BrCa patients stratified by a high vs low number of IR events. Samples have been dichotomised based on the median number of IR events.

Our analyses confirmed a previous report that BrCa is the only cancer in which the number of IR events is reduced compared to normal adjacent tissue (Figure 1B)^20^. All other cancers exhibit increased IR compared to their matched adjacent normal tissue (Figure 1B). However, we also noticed that the number of IR events in breast cancer itself was comparable with other cancers, while normal breast tissue presented with unusually high numbers of IR events (Figure 1B). In fact, normal breast tissue had the highest IR frequencies compared to all other normal tissues (Figure 1C). Therefore, reduced IR in tumours, which is unique to BrCa, can be largely attributed to normal breast tissue having the highest occurrence of IR events.

We applied beta regression models to identify differentially retained introns (dIRs) and found 3,024 dIRs between normal and breast cancer (Supplementary Figure 1A). Of these 210 were downregulated (in 160 genes) and 69 were upregulated (in 52 genes) with a ≥10% difference in the IR ratio (ΔIR ≥ 0.1). Downregulated IR events in BrCa are associated with processes related to cell cycle, nuclear division, and DNA replication among others (Supplementary Figure 1B).

Next, we explored whether the specific pattern of IR in BrCa is associated with clinical features. As shown in Supplementary Figure 2A and Figure 1D, IR patterns were distinct between BrCa vs normal tissues as well as estrogen receptors positive (ER^+^) vs negative (ER) samples, respectively. The human epidermal growth factor receptor 2 (HER2) amplified molecular subtype had the lowest average number of IR events (*n* = 1,731) compared to the other three main subtypes Luminal A (*n* = 2,089), Luminal B (*n* = 2,113), and Basal (*n* = 2,018) (Figure 1E). Strikingly, HER2-amplified tumours are associated with a 2.9-fold increased hazard ratio (*p* = 0.001, Supplementary Figure 2B). Moreover, advanced stage tumours (Stage III) had the lowest average number of IR events (*n* = 1,913) compared to Stage II (*n* = 2,064) and Stage I (*n* = 2,210) tumours (Figure 1F). Likewise, those tumours with the highest immunohistochemical (IHC) staining score for HER2 (score: 3) exhibited the lowest average number of IR events (*n* = 1,825) compared to score 1 (*n* = 2,095) or score 2 (*n* = 2,137) tumours (Supplementary Figure 2C). We also found that a high number of IR events is associated with better survival in patients with the Luminal B subtype (Figure 1G; Supplementary Figure 2D). Overall, these data support the conclusion that poor outcomes in BrCa correlate with low IR.

### Putative *trans*-regulators of IR in breast cancer

We confirmed that IR decreases in BrCa cells *in vitro* based on our analysis of cultured ER^+^ MCF7 cells and non-tumorigenic MCF10A cells (Figure 2A). Higher IR in a normal breast epithelial cell line (MCF10A) was associated with reduced gene expression (Supplementary Figure 3A).

**Figure 2.**
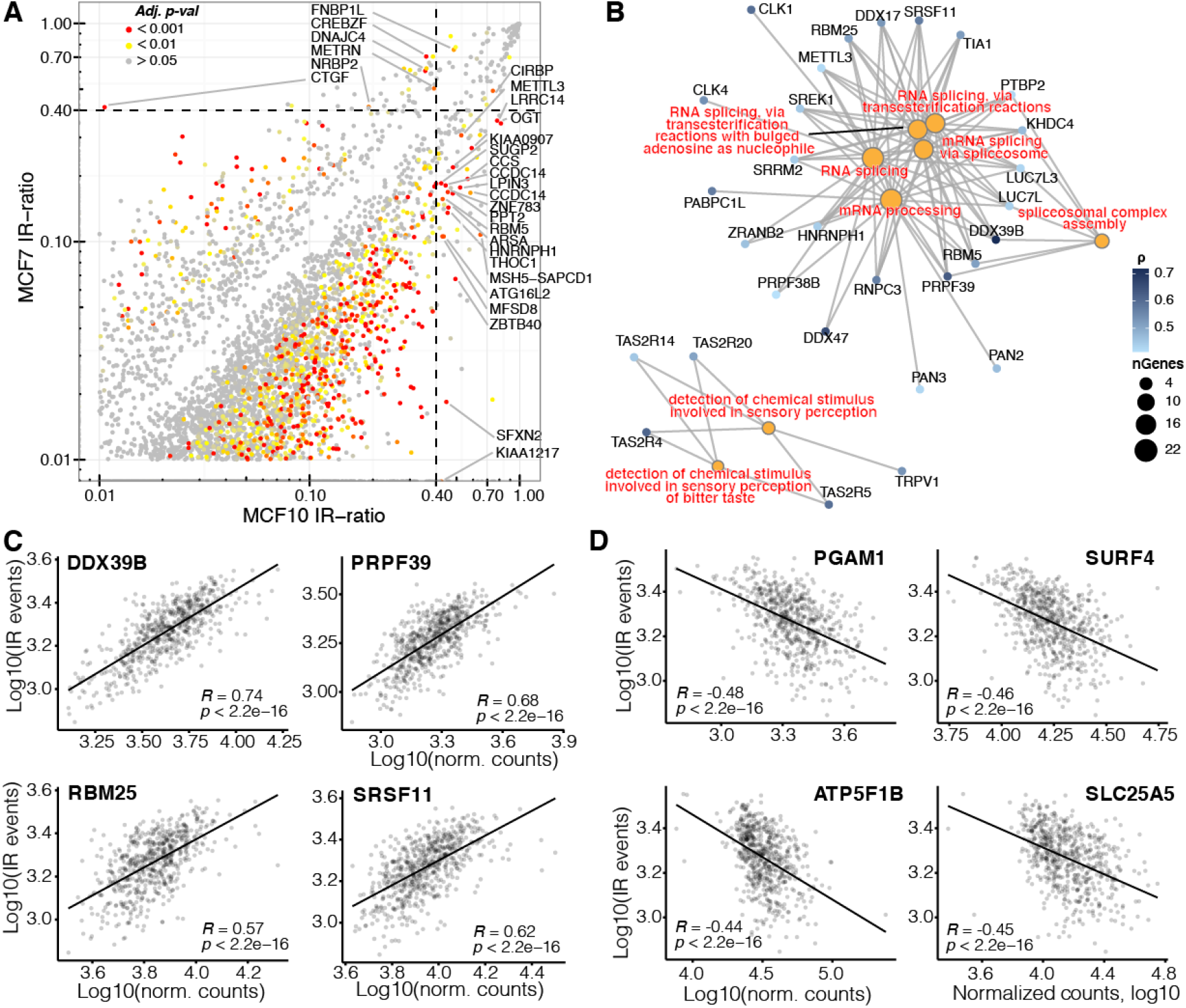
Putative regulators of intron retention in breast cancer. **(A)** Scatterplot showing differentially retained introns (dIR) between MCF7 and MCF10A cells. Genes associated with significant dIR events (p-adj. < 0.001) and IR-ratios > 0.4 are labelled. Given the low number of replicates (n=2) we have applied the Audic and Claverie test^21^ for significance testing. **(B)** Enriched GO terms in genes positively correlated with IR. Circle size of the GO terms (orange) is denoted by the number of genes associated with them. **(C)** RNA splicing associated genes that most highly correlate with the number of IR events in each sample. The scatterplots illustrate the log10 number of IR events against the log10 normalized read counts for each gene. **(D)** Genes that most strongly anti-correlate with the number of IR events in each sample.

To identify potential regulators of IR we correlated IR frequencies in the TCGA-BRCA cohort with expression values (normalized RNA-seq counts) of ~23,000 genes (Supplementary Table 2). We performed Gene Ontology (GO) enrichment analysis on the top 5% genes with the highest (*r* > 0.41) and lowest (*r* < −0.27) correlation coefficients to identify potential positive and negative regulators of IR, respectively. We found eight GO terms that were significantly enriched in positively correlated genes (*p-adj.* ≤ 0.05; Figure 2B). Intriguingly, six of these GO terms were related to RNA splicing, with *DDX39B*, *RBM25*, *PRPF39,* and *SRSF11* being among the genes that most highly correlate with the number of IR events in each sample (Figure 2C).

The four genes that most strongly anti-correlate with the number of IR events include the mutase PGAM1, the membrane protein encoding SURF4, the mitochondrial transmembrane transporter SLC25A5, and the mitochondrial ATP Synthase F1 Subunit Beta (ATP5F1B) (Figure 2D). Strikingly, the top 10 most significant GO terms (out of 399) associated with genes that negatively correlate with the number of IR events correspond to mitochondrial processes and cellular energetics (Supplementary Figure 3B).

### Highly proliferating cells have fewer IR events

Since the energy demand of a cell is tightly coupled with proliferation, we sought to examine a potential link between doubling times of BrCa cells and the number of IR events in 36 BrCa-related cell lines of the Cancer Cell Line Encyclopedia (CCLE). Indeed, we found that cells with slower doubling time exhibit a higher number of IR events (Figure 3A).

**Figure 3.**
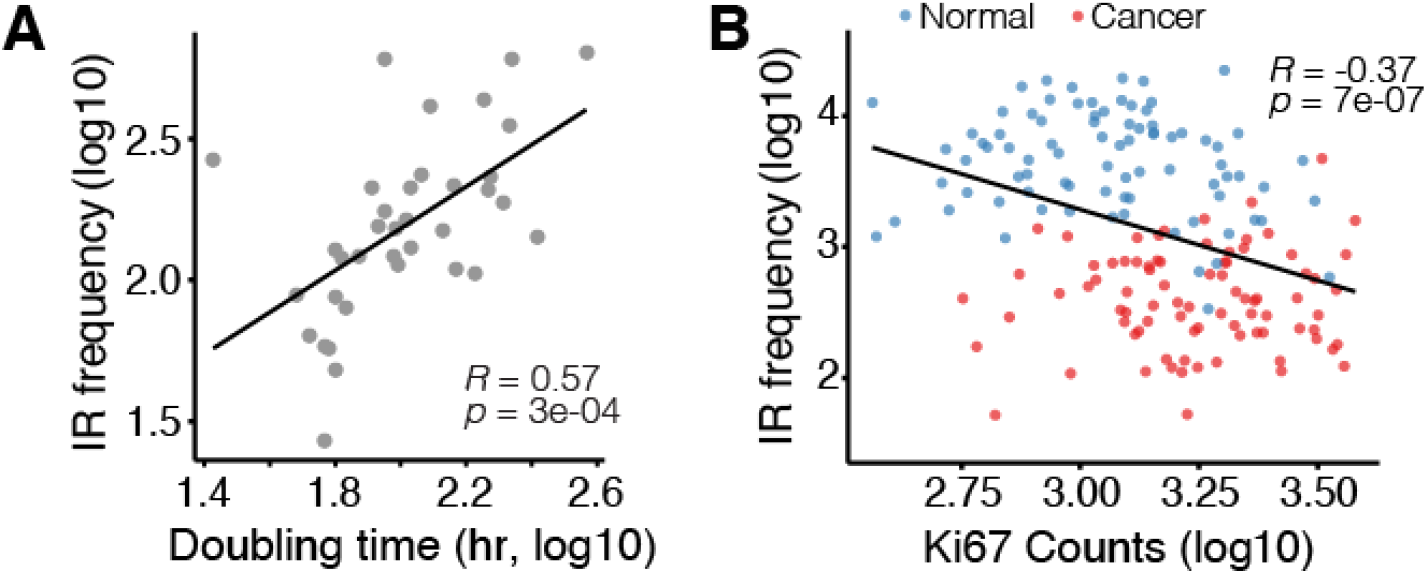
IR and cell proliferation. **(A)** Cancer Cell Line Encyclopedia (CCLE) cell doubling times (x-axis) correlate with number of IR events (y-axis). **(B)** Normalized read counts of proliferation marker Ki67 anti-correlate with the number of IR events (y-axis). Red dots – tumour samples; blue dots – normal breast tissue.

To corroborate this result, we also investigated whether a common proliferation marker would inversely correlate with IR frequencies. Since immunohistochemistry (IHC) staining of Ki-67 is unavailable for the TCGA cohort, we tested whether the proliferation rate in tissues might be estimated based on *MKI67* sequencing read counts. The *MKI67* gene encodes the proliferation marker protein Ki-67. Using Human Protein Atlas data, we confirmed that *MKI67* mRNA expression correlates with its Ki-67 protein staining intensities detected by IHC (Supplementary Figure 4A). As expected, normalised *MKI67* read counts were higher in all nine cancers when compared to the respective adjacent normal tissues (Supplementary Figure 4B). This suggests that *MKI67* read counts can be used as proxy for IHC staining to estimate cellular proliferation rates. *MKI67* expression was also found to be inversely correlated to the doubling time of 36 CCLE cell lines (Supplementary Figure 4C). We observed that the number of IR events in samples of the TCGA-BRCA cohort negatively correlated with the proliferation rate (Figure 3B).

### The role of RNA Binding Proteins in IR regulation

Next, we investigated genes that were specifically deregulated in BrCa. We identified a set of 150 genes that were *only* differentially expressed between BrCa and normal breast tissues, of which seven were RBPs (Figure 4A). We calculated the *z*-score of each gene’s log fold-change in BrCa versus the log fold-change in other cancers in order to estimate the level of specificity of a gene being differentially expressed in BrCa only (Figure 4B). Among the genes that are highly specifically over-expressed in BrCa are two known RBPs: *ZFP36L2* and *TUT4*. *ZFP36L2* promotes poly(A) tail removal of mRNA transcripts^22^, while Terminal Uridylyl Transferase 4 (*TUT4*) adds uridines to deadenylated transcripts^23^. Thus, both RBPs are mediators of mRNA decay, which could explain the observed reduction of IR transcripts in BrCa.

**Figure 4.**
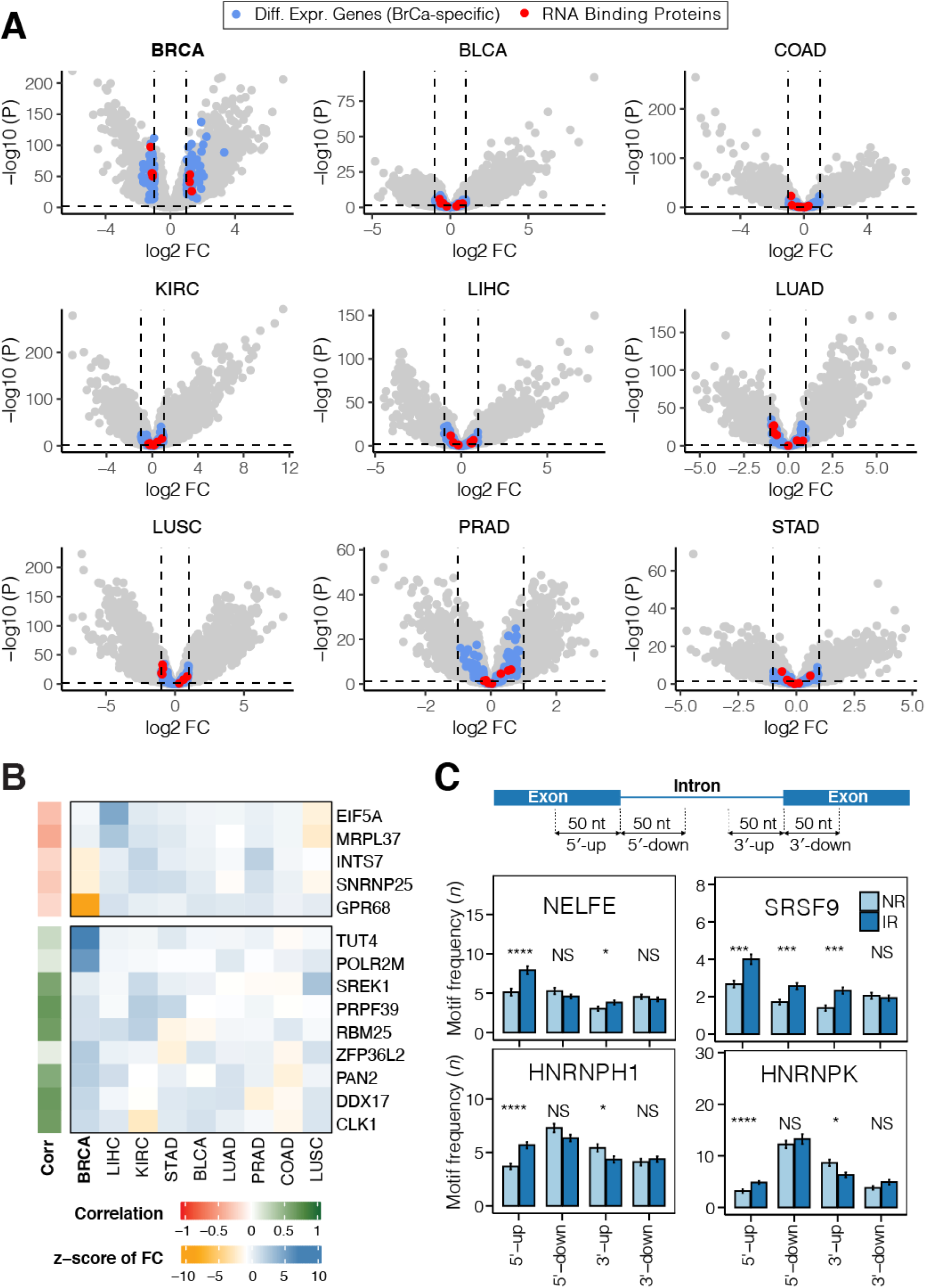
Breast cancer-specific gene expression and RBP analysis. **(A)** Volcano plots showing differentially expressed genes in nine tumour types vs adjacent healthy tissue. The dashed lines represent the *p*-value cut-off (horizontal; *p* < *0.05*) and fold-change threshold (vertical |*FC*| ≥ 1). See Figure 1A for cancer type abbreviations. Highlighted in blue are genes that are exclusively differentially expressed in BrCa, while those in red represent RBPs within this subset. **(B)** Heatmap of genes specifically differently expressed in BrCa (represented by colour-coded z-score). Annotation bar (left) shows the colour-coded correlation coefficient between gene expression and number of IR events in each sample. **(C)** Bar plots show the frequencies with which known binding motifs occur around the splice sites (50 nt up-/downstream) of differentially retained (IR; dark blue) and non-differentially retained introns (NR; light blue). Differences in average frequencies were determined using students *t*-test. * *p* < 0.05, *** *p* < 0.001, **** *p* < 0.0001, NS – not significant.

We also determined the frequency by which BrCa-specific genes occur in RNA-related gene sets (*n* = 138) in the Molecular Signatures Database (MSigDB; total ~5,000 curated gene sets)^24^. While known RBPs such as *ZFP36L2* and *SNRNP25* (part of the minor U12-type spliceosome) are annotated in multiple RNA-related gene sets, other genes, that are specifically differentially expressed in BrCa, did not show any potential RNA binding capabilities (Supplementary Figure 5A).

In addition, we analysed differentially retained introns for occurrences of RBP binding motifs. We found that differentially retained introns were enriched in NELFE and SRSF9 binding sites in upstream exons (5’-up) and the 3’ terminal region, respectively (Figure 4C). Moreover, retained introns have fewer HNRNPH1 and HNRNPK binding sites in their 3’ terminal region compared to non-retained introns (Figure 4C; Supplementary Figure 5B).

We conclude, that RPBs are among the factors that facilitate reduced IR in BrCa by enabling efficient splicing of introns from pre-mRNA transcripts. However, RBPs specifically differentially expressed in BrCa are not among those with enriched binding motifs within and around differentially retained introns. This suggests that more complex, multifaceted regulatory mechnisms are causing the consistent reduction of IR in BrCa.

### Tissue composition affects cancer IR profiles

Since the reduction in IR events in BrCa contrasts with all other cancer types analysed, we examined a possible contribution from the changing cell composition in the tumour microenvironment compared to healthy breast tissue. Gene signature-based and machine learning-based algorithms have been developed to deconvolute the cell type composition in bulk RNA-sequencing data^25^. To compare cell environmental profiles of TCGA breast tumour samples versus healthy adjacent control samples we used the cell type deconvolution algorithm xCell^26^, which was trained on 1,822 pure human cell type-specific transcriptomes extracted from single cell transcriptome profiling data. xCell analysis revealed that the breast tumour cell composition is distinct from other cancers (Supplementary Figure 6). Among the most enriched cell types in BrCa are T-helper cells, mesenchymal stem cells and basophils. These predictions are supported by recent single-cell BrCa profiling studies^27–29^. Normal breast tissue is enriched in endothelial cells, adipocytes, and dendritic cells (Figure 5A).

**Figure 5.**
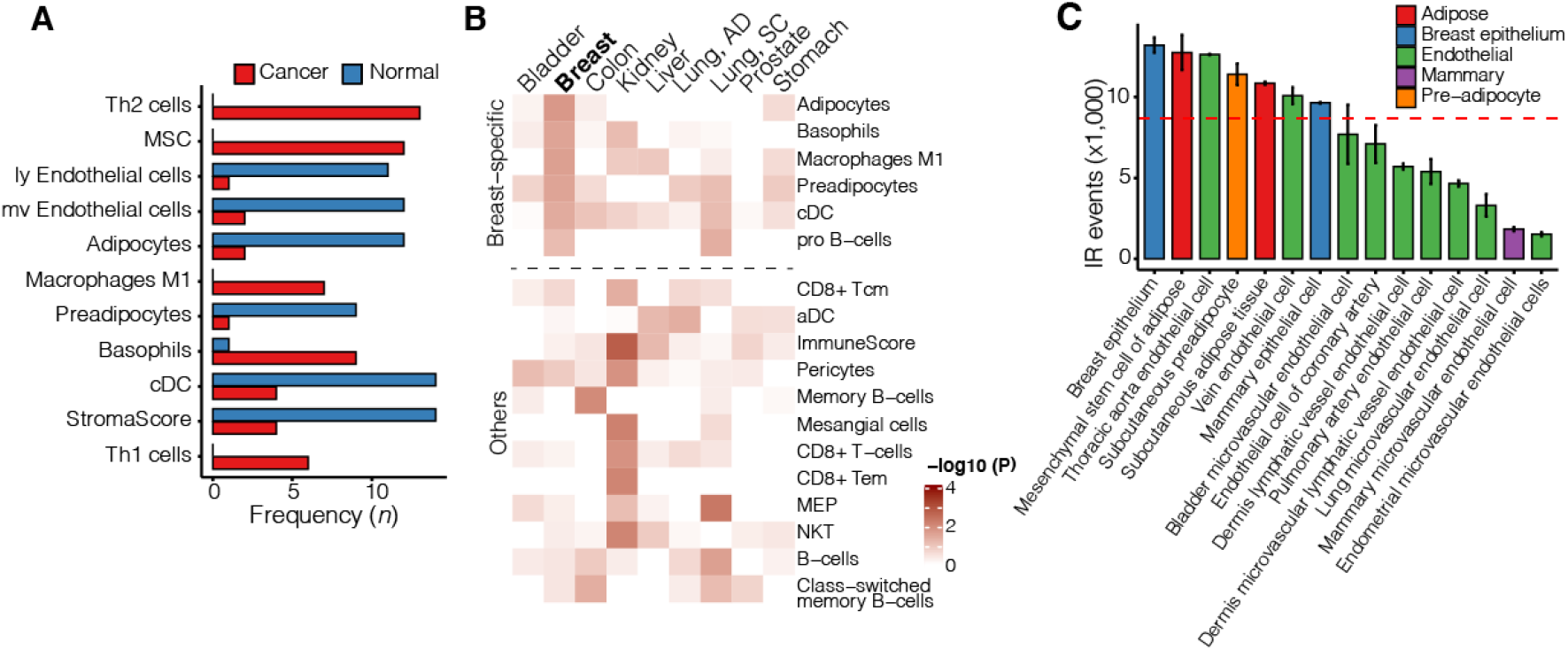
Breast tumour cell composition. **(A)** Frequently enriched cell types in breast tumours (red) and normal breast tissue (blue). **(B)** Heatmap illustrating cell type enrichment in healthy adjacent tissue of nine TCGA cancer cohorts. **(C)** Abundance of IR events in purified cells. Colours indicate groups of cells belonging to the same family. Dashed red line represent mean number of IR events. Th1/2 – T helper 1/2; MSC – mesenchymal stem cell; ly – lymphatic; mv – microvascular; a/cDC – activated/classical dendritic cell; Tcm – T central memory cell; Tem – T effector memory cell; NKT – natural killer T cell; MEP - Megakaryocyte– erythroid progenitor cell. ImmuneScore quantifies the enrichment of an immune cell signature including B-cells, T-cells, DC, eosinophils, macrophages, monocytes, mast cells, neutrophils, and NK cells. StromaScore quantifies the enrichment of a stroma-type cell signature including adipocytes, endothelial cells, and fibroblasts.

Indeed, adipocyte and myeloid cell (M1 macrophages, basophils) enrichment is specific to normal breast tissue (Figure 5B/C), which could explain the IR paradox in BrCa.

To determine whether cell types enriched in normal breast tissue have particularly high IR event frequencies we retrieved RNA sequencing data of 66 cell/tissue types from the ENCODE repository (Supplementary Table 3). Our analysis suggests that breast epithelial cells have the highest prevalence of IR followed by adipocytes (Figure 5C), which could explain the drop in IR events in breast tumours.

## Discussion

IR is omnipresent in vertebrate species^2,30^ and affects up to 80% of human protein-coding genes^17^. Numerous studies have highlighted the functional importance of retained introns in a wide range of biological functions including cell differentiation and development ^12,31–34^.

Since first reports in 2015 and subsequent confirmatory studies, BrCa has stood in stark contrast to other cancers in relation to its burden of IR^20^. Dysregulation of *cis*- and *trans*- modulators can cause aberrant IR in various cancers^20^. For example, Dvinge *et al.* found that snRNA expression changes IR in the MCF7 cell line and to a certain degree in BrCa patient samples. They also showed that splicing factor knockdown can lead to increased IR in triple-negative BrCa (TNBC)^35^. Kim *et al.* found that some BrCa IR events anti-correlate with DNA methylation and that high IR levels in transcripts of migration and invasion inhibitory protein (MIIP) are associated with increased survival in European-American patients with invasive breast carcinoma^36^.

We confirmed a consistent reduction of IR events in TCGA breast adenocarcinoma samples compared to adjacent normal breast tissue. While BrCa is the only cancer where this reduction is observed, IR frequencies are, in fact, comparable to those observed in other cancer types. This is due to the excessively large number of IR events in healthy breast tissue. Gascard *et al.* found that IR increases with differentiation state in normal human breast cells with fewer IR events in myoepithelial cells and seven times more events in luminal epithelial cells^37^. Indeed, our results suggest that an important factor in the reduction of IR events in breast tumours is the changing cell composition from adipocyte and epithelial cell-rich breast tissue to lymphocyte-infiltrated breast tumours. Adipocytes and epithelial cells have one of the highest IR frequencies in their transcriptomes compared to other cell types, while lymphocytes are known to have low IR counts^38^. Siang and co-workers have shown in this context, that the RBP human antigen R (HuR), which is involved in pre-mRNA processing, is a negative regulator of adipogenesis^39^. Interestingly, Diaz-Muñoz *et al.* demonstrated that HuR binding to introns modulates alternative intron usage^40^. This may contribute to the high IR observed in adipocyte-rich normal breast tissue.

Aberrant IR has previously been associated with disease phenotypes and clinical outcomes. For example, IR in *CMYC* and *SESTRIN1* genes was shown to be a reliable molecular marker separating melanoma from non-melanoma tumours^14^ and Sznajder and colleagues have shown that IR can be used as biomarker in hereditary repeat expansion diseases^15^. Despite marked differences between tumour and normal breast tissue, IR profiles in our analysis also differ between ER^+^ vs ER^−^ tumours. The survival advantages associated with high IR numbers in the Luminal B subtype suggest that this form of alternative splicing should be considered for therapeutic exploitation. However, the exact mechanisms whereby dynamic IR profiles lead to differences in clinical outcomes would be the subject of future studies.

The inverse relationship between IR and cell proliferation has been previously observed in the context of B-cell development and T-cell activation^38,41^. Our results demonstrate that the number of IR events positively correlates with longer cancer cell doubling times and that more IR events are associated with slower cell proliferation in BrCa. Our data shows that HER2 positive breast tumours have the lowest number of IR events. HER2 is known to induce cell proliferation in human cancers and is associated with poor prognosis in BrCa^42^. These results suggest that IR is a mechanism that counteracts tumour growth and would provide opportunities as therapeutic targets. Interestingly, the tumour suppressor Herstatin, expressed in healthy breast tissue^43^, is a splice variant of the oncogene *HER2*, with a retained intron 8^44^. Herstatin is a secreted autoinhibitor of Her2^44^ and intron 8 retention is regulated by RBPs of the HNRNP1 family (including H1, D, and A2/B1)^45^. Koedoot and co-workers have demonstrated that inhibition of cell proliferation can be achieved via splicing factor knockdown in TNBC^46^.

In summary, our study sheds light on the unique causes and consequences of aberrant splicing in BrCa. The modulation of IR levels may offer novel opportunities for personalised BrCa treatment, especially in hormone- and chemotherapy-resistant subtypes.

## Materials and methods

### RNA-sequencing data/patient samples

We retrieved data from nine tumour types and healthy adjacent tissue, including 615 BrCa patient samples generated by the TCGA (Figure 1A). Only samples for which sequencing had been performed at >40M read depth were selected for analysis. Moreover, only tumour types with at least 20 matched tumour/normal tissue samples where considered.

RNA-seq data were downloaded as BAM files using the R/Bioconductor package TCGAbiolinks^47^ and the command-line tool gdc-client v1.4.0 (github.com/NCI-GDC/gdc-client) under an approved data access application. All files were checked for integrity. Harmonized gene expression data in the form of HTseq counts^48^ were downloaded using TCGAbiolinks^47^.

### mRNA-sequencing and data analysis – MCF7 and MCF10A cells

Total RNA was isolated from MCF7 and MCF10A cells using Trizol (Invitrogen). The RNA quality was assessed using RNA 6000 Nano Chips on an Agilent Bioanalyzer (Agilent Technologies) to confirm an RNA integrity score of >7.0. mRNA-seq was performed by Macrogen (Korea) using the Illumina Hi-Seq 2000 platform. RNA-seq libraries were prepared from > 1 μg of total RNA using TruSeq RNA sample prep kit (Illumina) according to the manufacturers’ instructions.

### Differential IR and gene expression analyses

IR was quantified using IRFinder v1.2.0 ^17^, using the Ensembl human genome (hg38, release 86) as reference. The IRFinder algorithm measures 20 parameters for IR detection in each sample, including the median number of reads mapping to each nucleotide across the intron length (intron depth, ID), the ratio of nucleotides within an intron with mapped reads (coverage), the number of reads that map to the 5′ flanking exon and to another exon within the same gene (splice left, SL), the number of reads that map to the 3′ flanking exon and to another exon within the same gene (splice right, SR), the number of reads spanning the exon-exon junction (splice exact, SE) as well as the IR ratio:

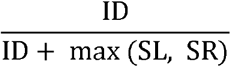

The following criteria were used for quantifying the number of IR events in a sample:

1. 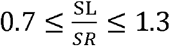;
2. (*SL* + *SR*) > 10 in ≥50% of samples;
3. *coverage* > 0.5 in ≥50% of samples;
4. IR > 0.05 in at ≥50% normal or cancer samples

The number of IR events in a sample was determined based on introns with an IR ratio > 0.1 and meeting the filtering criteria described above. Introns not meeting these criteria were not considered as being retained. Beta regression was used to identify differentially retained introns (dIR) between cancer and adjacent normal tissues using the betareg R package ^49^. Since IR ratios are proportional data with values between 0 and 1, we reasoned that beta regression was best suited to model IR and identify dIRs between normal and cancer tissues. An absolute difference in the IR ratio (ΔIR = IR_Cancer_ − IR_Nornal_) of more than 0.1 with FDR-adjusted *p* < 0.05 was considered significant.

Dimensionality reduction, i.e. principal component analysis (PCA), of IR profiles was performed using the package factoextra (github.com/kassambara/factoextra).

Differential gene expression between normal breast tissue and BrCa was performed using the DESeq2 package^50^. Genes with an average read count >10 in all samples were selected for differential gene expression analysis (*n* = 23,072). Genes with an absolute log2 fold-change > 1 and FDR-adjusted *p* < 0.05 were considered significant. To identify genes that were specifically differentially expressed in BrCa, we removed genes that were differentially expressed in any of the other 8 cancers and determined specificity by computing the z-score on log-fold change using the log-fold change observed in BrCa as reference.

### Gene Ontology and RBP analyses

Gene Ontology analysis was performed using the clusterProfiler package^51^. The false discovery rate (FDR) approach was used for multiple testing correction. The list of 1,542 RBPs was taken from Gerstberger *et al.*^52^.

### Survival analysis

Patient survival data was provided by the TCGA consortium. Survival analysis was performed using packages Surv and survminer (github.com/kassambara/survminer).

### RNA binding protein motif detection

RNA binding protein (RBP) motifs in position-weight matrix format (PWM) were retrieved from the ATtRACT database (version 0.99*β*)^53^, which contains 1,196 motifs corresponding to 160 human RBPs. Sequences of 100 nt were extracted from the regions flanking retained and non-retained introns and scanned for the presence of motifs using the fimo tool provided by the meme suite^54^.

## Acknowledgements

The results shown here are based upon data generated by the TCGA Research Network: https://www.cancer.gov/tcga. The authors acknowledge The University of Sydney High Performance Computing service at The University of Sydney for providing resources that have contributed to the research data reported within this paper.

Financial support was provided by National Health & Medical Research Council Investigator Grants (#1177305, #1196405) to J.E.J.R. and U.S. and Project Grants (#507776, #1128748) to J.E.J.R. We also received support from the Cancer Council NSW (project grants RG11-12, RG14-09, RG20-12 to J.E.J.R. and U.S.). M.M. is funded through a Victoria Cancer Agency Fellowship.

## Authors’ contributions

J.S.S. and U.S. designed the research. J.S.S., U.S., and V.P. performed bioinformatic analyses and interpreted the data. A.Y.M.A., J.J.L.W., and J.E.J.R. performed and/or supervised experiments. J.S.S. and U.S. wrote the manuscript with help from M.M., J.E.V., J.J.L.W., and J.E.J.R. All authors have read and agreed to the published version of the manuscript.

