## Supplementary Materials for "Towards resolution of the intron retention paradox in breast cancer"

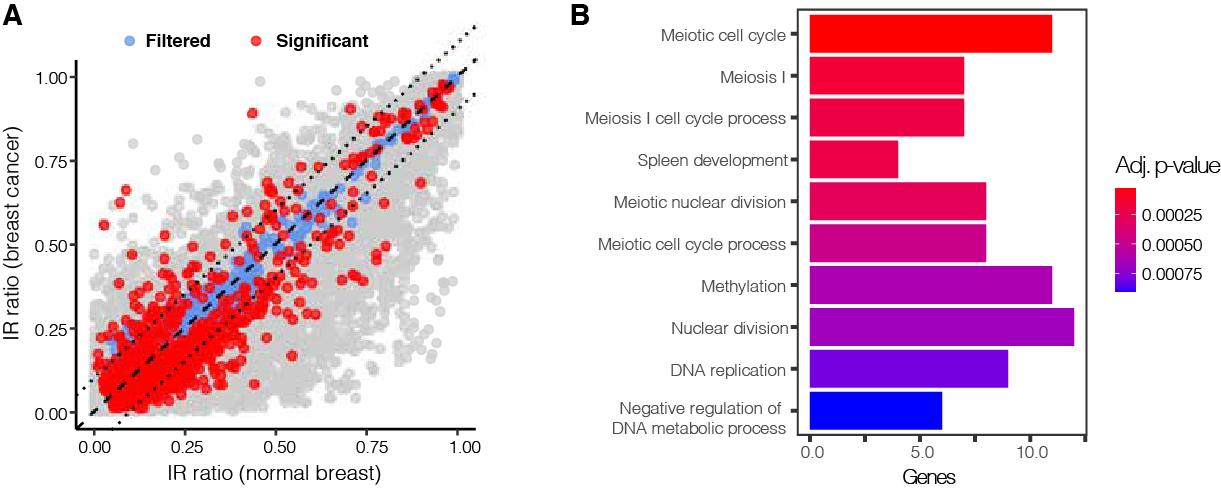

Supplementary Figure 1 Intron retention in breast cancer versus normal adjacent tissue. (A) IR ratios of differentially retained introns in BrCa and normal breast tissue (blue – filtered introns; red – significantly differentially retained introns). (B) Gene Ontology enrichment of genes with reduced IR in BrCa. The analysis is based on n = 150 matched samples.

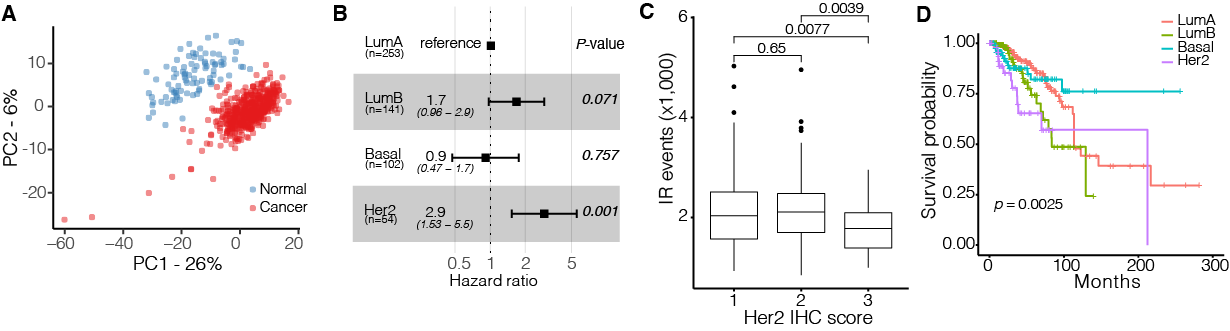

Supplementary Figure 2 Clinical relevance of IR in breast cancer. (A) Principal component analysis (PCA) plot illustrating distinct IR profiles of BrCa (red) and normal breast tissue (blue) samples (*n* = 615). (B) Cox hazard ratio for BrCa subtypes. Luminal A breast cancer was chosen as reference as it is the most benign subtype of BrCa and exhibits the greatest level of IR. (C) Distributions of IR event frequencies in tumour samples were assigned to each of the three HER2 immunohistochemistry (IHC) scores. (D) Kaplan-Meier plot showing the survival probability of BrCa patients with different molecular subtypes.

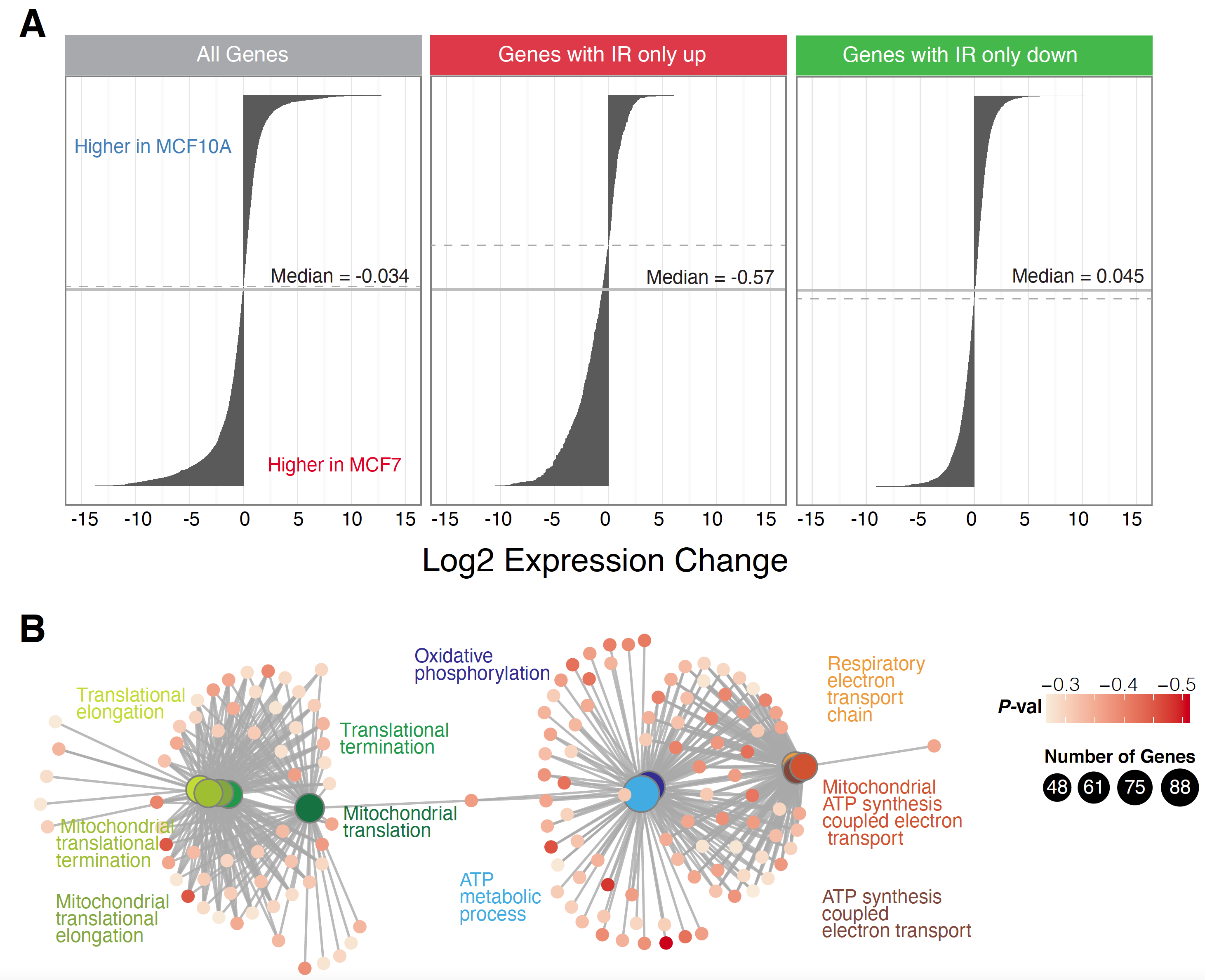

Supplementary Figure 3 Intron retention effects on gene expression. (A) Waterfall plots showing the distribution of all up- and downregulated genes (left), those that have increased (middle) or decreased (right) IR levels in MCF10a vs MCF7. Dashed lines indicate inflection point. (B) Top 10 most significant GO terms associated with genes that negatively correlated with the number of IR events.

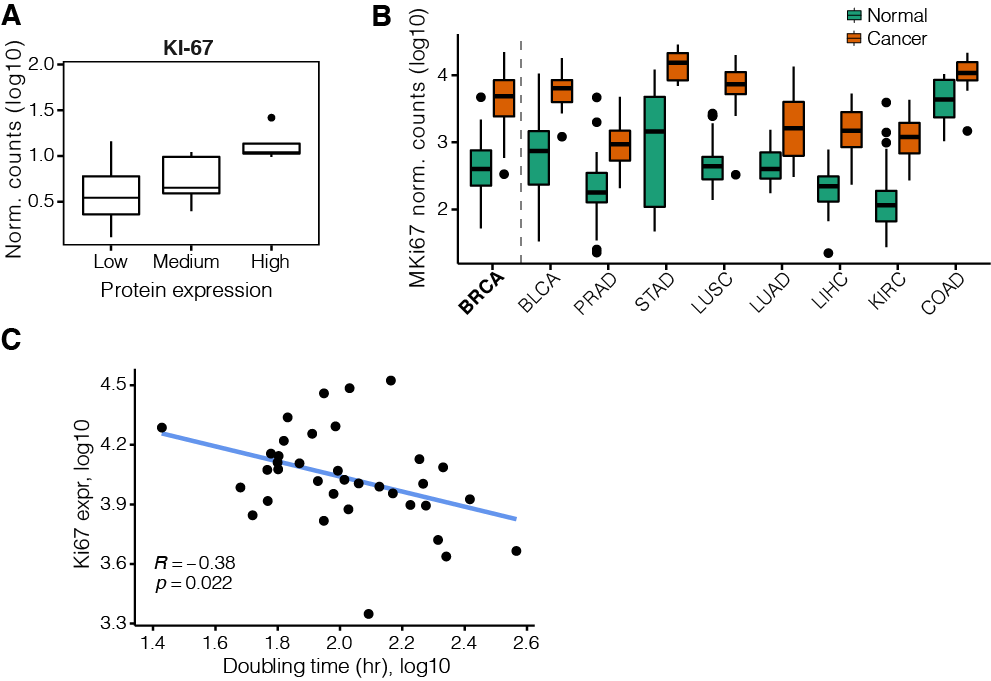

Supplementary Figure 4 Proliferation markers of cancer cells. (A) *MKI67* mRNA expression corresponds to Ki-67 protein levels based on immunohistochemistry data (Human Protein Atlas). Box-whisker plots represent log10 normalised counts of *MKI67* and *PCNA,* two cell proliferation marker genes. Horizontal lines indicate median count number. Dots represent outliers. (B) Normalized *Ki67* read counts used as proliferation index for nine TCGA tumour types. Dots represent outliers. (C) Anti-correlation of *MKI67* expression (normalised read counts; y-axis) and cell doubling time (x-axis) in CCLE BrCa cell lines.

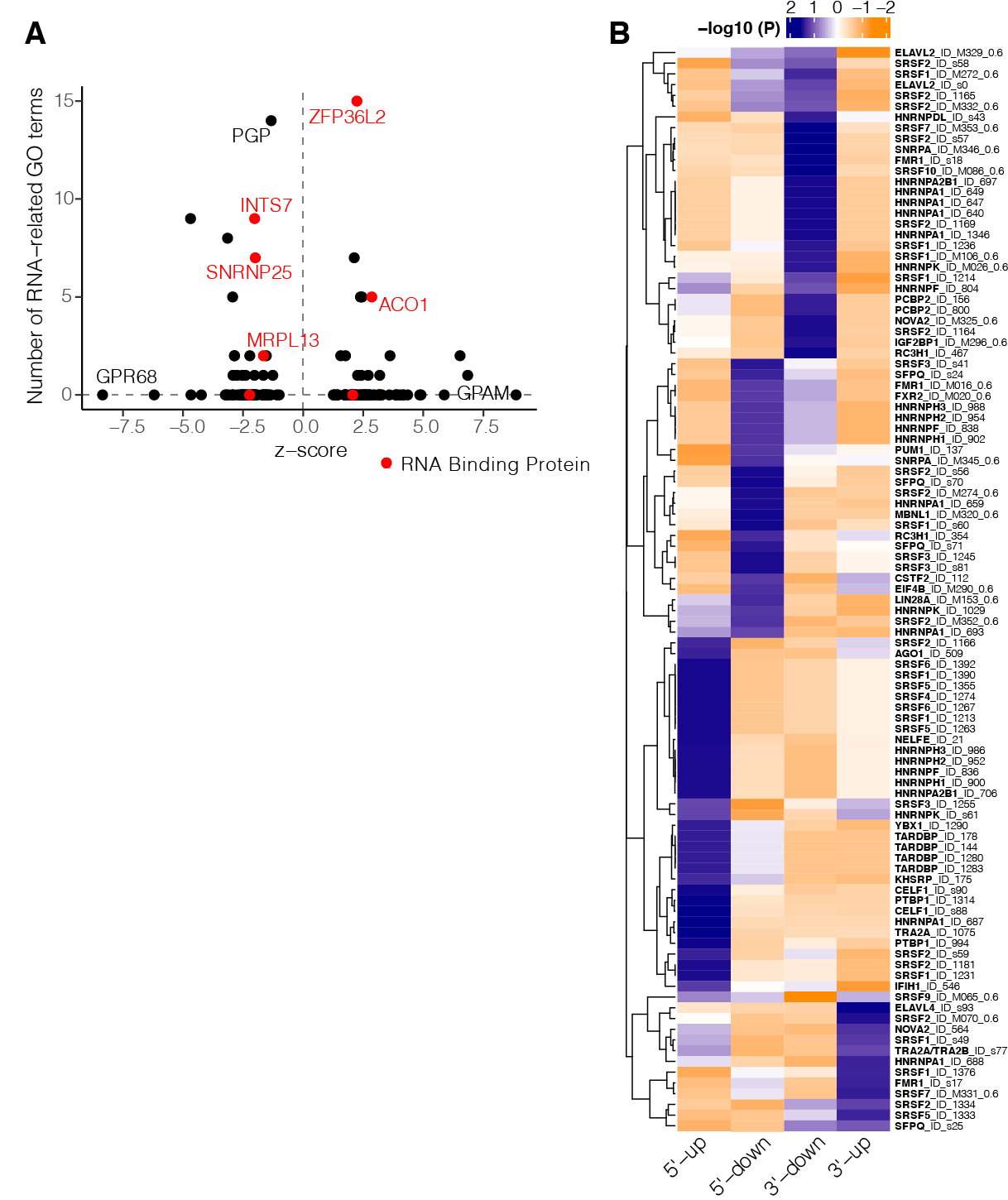

Supplementary Figure 5 RBP motif enrichment. (A) Frequency of BrCa-specific genes occuring in RNA-related gene sets (n = 138) in the Molecular Signatures Database. Highlighted in red are RNA binding proteins. (B) Heatmap of enriched RBP binding motifs.

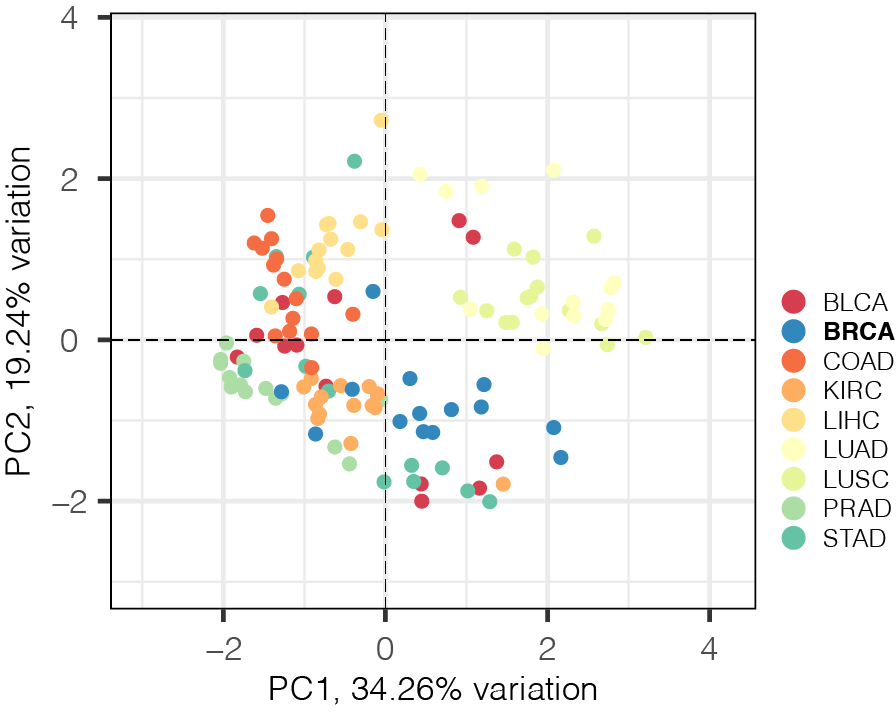

Supplementary Figure 6 Principal component analysis of tumour cell composition profiles.
